# Cohesion: A method for quantifying the connectivity of microbial communities

**DOI:** 10.1101/112391

**Authors:** Cristina M. Herren, Katherine D. McMahon

## Abstract

The ability to predict microbial community dynamics lags behind the quantity of data available in these systems. Most predictive models use only environmental parameters, although a long history of ecological literature suggests that community complexity should also be an informative parameter. Thus, we hypothesize that incorporating information about a community’s complexity might improve predictive power in microbial models. Here, we present a new metric, called community “cohesion,” that quantifies the degree of connectivity of a microbial community. We validate our approach using long-term (10+ year) phytoplankton datasets, where absolute abundance counts are available. As a case study of our metrics’ utility, we show that community cohesion is a strong predictor of Bray-Curtis dissimilarity (R^2^ = 0.47) between phytoplankton communities in Lake Mendota, WI, USA. Our cohesion metrics outperform a model built using all available environmental data collected during a long-term sampling program. The result that cohesion corresponds strongly to Bray-Curtis dissimilarity is consistent across the five lakes analyzed here. Our cohesion metrics can be used as a predictor for many community-level properties, such as phylogenetic diversity, nutrient fluxes, or ecosystem services. We explain here the calculation of our cohesion metrics and their potential uses in microbial ecology.

**Conflict of Interest:** The authors declare no conflict of interest.

## Introduction

Most efforts to model microbial communities primarily use environmental drivers as predictors of community dynamics (Patterson 2009, Hambright et al. 2015). However, despite the vast quantities of data becoming available about microbial communities, predictive power in microbial models is often surprisingly poor (Blaser et al. 2016). Even in one of most well-studied microbial systems, the San Pedro Ocean Time Series (SPOT), there are sampling sites where none of the 33 environmental variables measured are highly significant (P < 0.01) predictors of community similarity (Cram et al. 2015). Thus, there may be room to improve predictive models by adding new parameters; ecological literature has long suggested that the degree of complexity in a community should inform community dynamics (MacArthur 1955, Cohen and Newman. 1985, Wootton and Stouffer 2016). We hypothesize that incorporating information about the complexity of microbial communities could improve predictive power in these communities.

Here, we present a workflow to generate metrics quantifying the connectivity of a microbial community, which we call “cohesion.” We show how cohesion can be used to predict community dynamics (e.g. rate of compositional turnover, phylogenetic diversity, ecosystem services provided by the community). As an application of our metrics, we present a case study using our newly developed cohesion variables as predictors of the compositional turnover rate (a common response variable in microbial ecology) in phytoplankton communities.

Our cohesion metrics overcome two barriers that often preclude using information about community complexity in microbial analyses. First, the large number of taxa in microbial datasets makes it difficult to use information about all taxa in statistical analyses. Although methods exist to analyze microbial community interconnectedness (e.g. Local Similarity Analysis, artificial neural networks), this often involves constructing networks with many parameters that are difficult to interpret. Second, microbial community data are often “relativized” or “compositional” datasets, where the abundance of each taxon represents the fraction of the community it comprises. This creates several problems in downstream analysis (Weiss et al. 2016). For example, taxon correlation values are different in absolute versus relative datasets (Faust and Raes 2012, Friedman and Alm 2012), and it is unclear how using relative abundances influences the apparent population dynamics of individual taxa (Lovell et al. 2015). Thus, these two features (many taxa and relative abundance) have previously proven problematic when analyzing microbial community connectivity.

Here, we describe and test a method to quantify one aspect of microbial community complexity. Our resulting “cohesion” metrics quantify the connectivity of each sampled community. Thus, our cohesion metrics integrate easily with other statistical analyses and can be used by any microbial ecologist interested in asking whether community interconnectedness is important in their study system. We demonstrate how to obtain these cohesion metrics from time series data and, as a case study, show how cohesion relates to rates of compositional turnover in long-term phytoplankton datasets. We develop this workflow with datasets where raw abundance data are available and use these raw abundances to validate our methods when working with relativized datasets. Thus, our approach was designed to overcome known challenges of analyzing microbial datasets.

## Methods and Results

### Description of datasets

We restricted our search for datasets to those measured in absolute abundance that spanned multiple years, as to cover a wide range of environmental conditions. We found several datasets hosted on the North Temperate Lakes Long Term Ecological Research (NTL LTER) database that were over 10 years in length and contained counts of phytoplankton in absolute abundance. The datasets are from the following lakes in Wisconsin, USA: Lake Mendota (293 samples with 410 taxa over 19 years), Lake Monona (264 samples with 382 taxa over 19 years), Paul Lake (197 samples with 209 taxa over 12 years), Peter Lake (197 samples with 237 taxa over 12 years), and Tuesday Lake (115 samples with 121 taxa over 12 years). These lakes vary in size, productivity, and food web structure. Peter Lake and Tuesday Lake were also subjected to whole-lake food web manipulations during the sampling timeframe (detailed in Elser and Carpenter 1988 and Cottingham et al. 1998). We present the workflow using results from the Lake Mendota dataset, as it is the dataset of the longest duration with the largest number of taxa. Details about datasets can be found at https://lter.limnology.wisc.edu/.

### Data curation

Phytoplankton densities in Lake Mendota varied by more than 2 orders of magnitude between sample dates. Densities of cells in these samples ranged from 956 cells/mL to 272 281 cells/mL. We removed individuals that were not identified at any level (e.g. categorized as Miscellaneous). For each sample date, we converted the raw abundances to relative abundances, such that all rows summed to 1. We removed taxa that were not present in at least 5% of samples, as we were not confident that we could recover robust connectedness estimates for very rare taxa. This cutoff retained an average of 98.9% of the identified cells in each sample. Results of our analyses using other cutoff values can be found in the SOM.

### Overview

The input of our workflow is the taxon relative abundance table, and the outputs are measurements of the connectivity of each sampled community, which we call community “cohesion” (Fig. 1). In the process, our workflow also produces metrics of the connectedness of each taxon. Briefly, our workflow begins by calculating the pairwise correlation matrix between taxa, using all samples. We use a null model to account for bias in these correlations due to the skewed distribution of taxon abundances (i.e. many small values and a few large values) and relativized nature of the dataset (i.e. all rows sum to 1). We subtract off these “expected” correlations generated from the null model to obtain a matrix of corrected correlations. For each taxon, the average positive corrected correlation and average negative corrected correlation are recorded as the connectedness values. As previously noted, our goal was to create a metric of connectivity for each community; thus, the next step in the workflow calculates cohesion values for each sample. Cohesion is calculated by multiplying the abundance of each taxon in a sample by its associated connectedness values, then summing the products of all taxa in a sample. There are two metrics of cohesion, because we separately calculate metrics based on the positive and negative relationships between taxa. Within each section (1, 2, and 3), we alternate between presenting an analysis step and showing a validation of these techniques.

**Figure 1:**
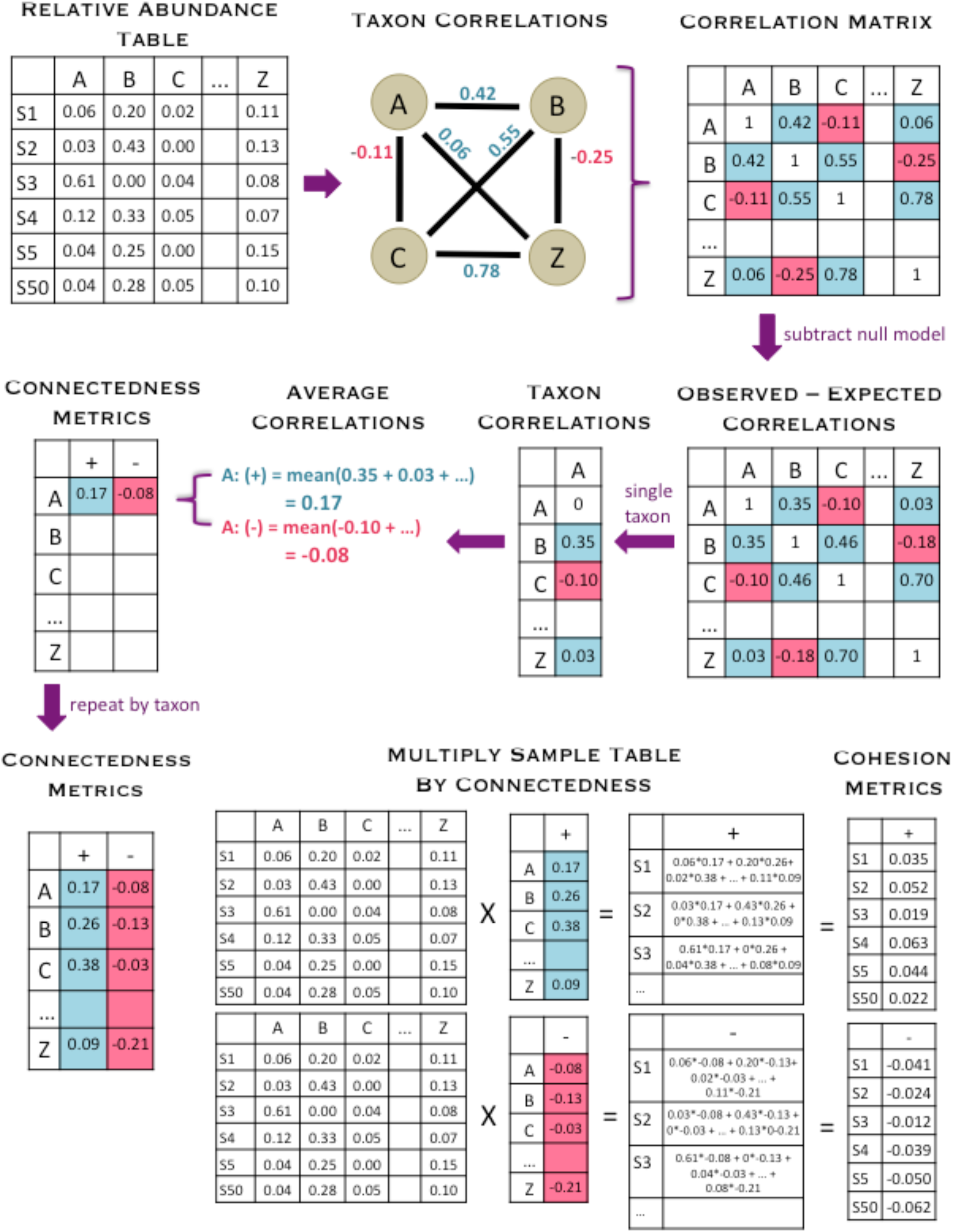
This diagram shows an overview of how our cohesion metrics are calculated, beginning with the relative abundance table and ending with the cohesion values. The relative abundance table shows six samples (S1 indicating “Sample 1”, etc.) and a subset of taxa (A, B, C, and Z). First, pairwise correlations are calculated between all taxa, which are entered into the correlation matrix. We then used a null model to account for how the features of microbial datasets might affect correlations, and we subtracted off these values (null model detailed in Fig. 2). For each taxon, we averaged the positive and negative corrected correlations separately and recorded these values as the positive and negative connectedness values. Cohesion values were obtained by multiplying the relative abundance table by the connectedness values. Thus, there are two metrics of cohesion, corresponding to positive and negative values.

#### 1. CONNECTEDNESS METRIC

##### Analysis

It is difficult to directly observe interactions within microbial communities, so correlations are often used to infer relationships between taxa or between a taxon and the environment. Thus, we used a correlation-based approach for determining the connectedness of taxa. However, when using correlation-based approaches with relativized microbial datasets, it is necessary to use a null model to evaluate how the features of the dataset (skewed abundances and the fact that all rows sum to 1) contribute to correlations between taxa (Weiss et al. 2016). The purpose of a null model is to assess the expected strengths of correlations when there are no true relationships between taxa (Ulrich and Gotelli 2010).

Our null model was used to calculate how strongly the features common to microbial datasets contribute to taxon connectedness estimates, so that this structural effect could be subtracted from the connectedness metrics. While testing various null models, it became clear that a taxon’s pairwise correlation values were strongly related to its mean abundance and persistence (fraction of samples when present) across the dataset. Thus, these features (mean relative abundance and persistence) were retained in the null model by only randomizing abundance values within each taxon. Details about alternate null models investigated are given in the SOM.

The objective of the null model was to calculate the strength of pairwise correlations that would be observed if there were no true relationship between taxa. During each iteration, one taxon was designated as the “focal taxon” (Fig. 2). For each taxon *besides the focal taxon*, abundances in the null matrix were randomly sampled without replacement from their abundance distribution across all samples. Then, we calculated Pearson correlations between the focal taxon and the randomized other taxa. We iterated through this process of calculating pairwise correlations between the focal taxon and all other taxa 200 times. The median correlations from these 200 randomizations were called the “expected” correlations for the focal taxon. We repeated this process for each taxon as the focal taxon, which resulted in a matrix of expected taxon correlations. Finally, we subtracted the expected taxon correlations from their paired observed taxon correlations, thereby producing a matrix where each value was an observed minus expected correlation for the given pair of taxa.

**Figure 2:**
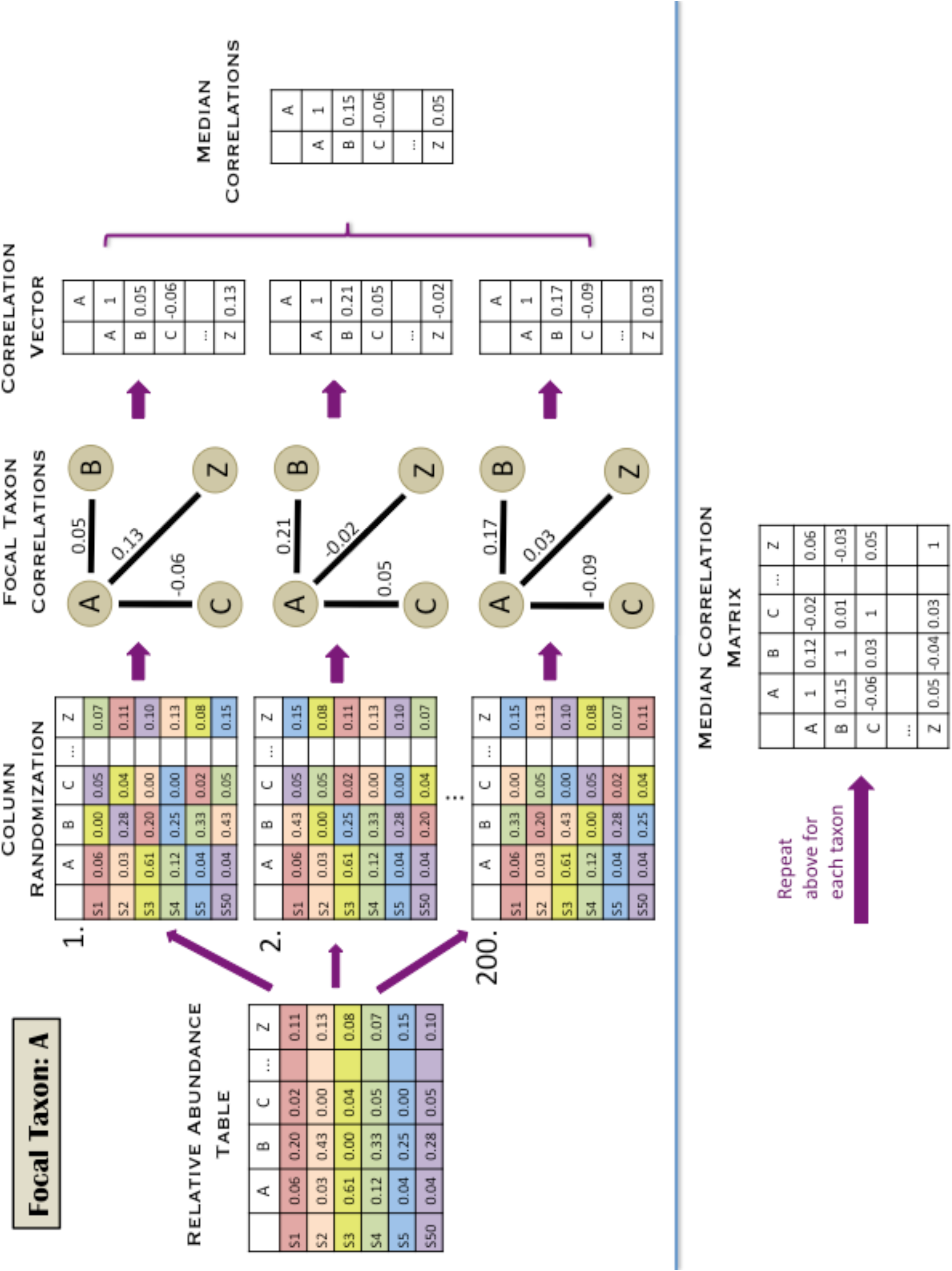
Microbial data are in the form of relative abundance, and some taxa are much more abundant than others, which are factors that may cause taxa to be spuriously correlated. Thus, we devised a null model to account for the bias that these data features introduce into our metrics. We repeated this process with each taxon as the “focal taxon,” which is A in this figure. For each of 200 iterations, we randomized all taxon abundances *besides the focal taxon*. We then calculated correlations between the focal taxon and all other taxa. We recorded the median value of the 200 correlations calculated for each pair of taxa in the median correlation matrix.

We calculated taxon connectedness values from the corrected (observed minus expected) correlation matrix. For each taxon, we separately averaged its positive and negative correlations with other taxa to produce a value of positive connectedness and a value of negative connectedness. We kept positive and negative values separate, because converting the dataset to relative abundance should affect positive and negative correlations differently, since an increase in the relative abundance in one taxon must result in the decrease in the relative abundances of other taxa.

The averaging step in this workflow was intended to overcome the issue that individual correlations between taxa can be influenced by many factors and may be spurious (Fisher and Mehta 2014). However, assuming that correlations often (but not always) reflect complexity in a community, the average of many correlations should be a more robust metric of complexity than any single correlation. In other words, we assume only that highly connected taxa have stronger correlations *on average*. Invoking the law of large numbers, these average correlations should be increasingly accurate measures of a taxon’s connectedness as the number of pairwise correlations increases (i.e. as the number of taxa in the dataset increases).

##### Validation

As discussed previously, there are inherent limitations of using correlation-based methods with relative abundance data instead of absolute counts (Fisher and Mehta 2014). Thus, we examined whether a measure of connectedness based on absolute abundance would show the same pattern observed using the relativized data. However, we needed a different approach for calculating correlations in order to account for the following properties of count data: 1) variance-mean scaling, which results in very large population variances of abundant taxa (Taylor 1961) and 2) the fact that individual population sizes are strongly related to overall community densities, which causes positive correlations among all taxa (Doak et al. 1998). As noted previously, phytoplankton densities in Lake Mendota samples varied by more than 2 orders of magnitude among sample dates. Therefore, using correlations between raw abundances would inflate the positive relationships between taxa as a result of changing overall community density. Thus, we first detrended the count data to account for changing community density (on different sampling dates) and drastically different variances of taxon populations (which are expected as a result of mean-variance scaling).

We used a hierarchical linear model to estimate the effects of overall community density and mean taxon abundance on individual taxon observations (sensu Jackson et al. 2012), so that these effects could be removed when calculating correlations. We modeled the abundance of each taxon at each time point as a function of sample date and taxon, assuming a quasipoisson distribution (which accounts for increases in population variances when population means increase). The residuals of this analysis represent the normalized (transformed) deviations of taxon abundances after accounting for overall community density on the sample date and taxon abundance/variance. We created a pairwise correlation matrix for the phtoplankton taxa using the correlations between these residuals. We calculated connectedness metrics from the pairwise correlation matrix using the same technique that we applied to the corrected correlation matrix from the relativized data: we used the average positive and negative taxon correlations as their connectedness values.

We validated our workflow for the relative abundance dataset using the estimates of taxon connectedness obtained from the absolute abundance dataset. Comparing the connectedness values from these two methods shows strong agreement between the two sets of connectedness metrics (correlation for positive connectedness metrics = 0.820; correlation for negative connectedness metrics = 0.741, Fig. 3). Although two taxa deviate from the linear relationship between the negative connectedness metrics (appearing as outliers in Fig. 3B), both metrics rate these taxa as having strong connectedness arising from negative correlations. Thus, the two methods are qualitatively consistent for these two anomalous points.

**Figure 3:**
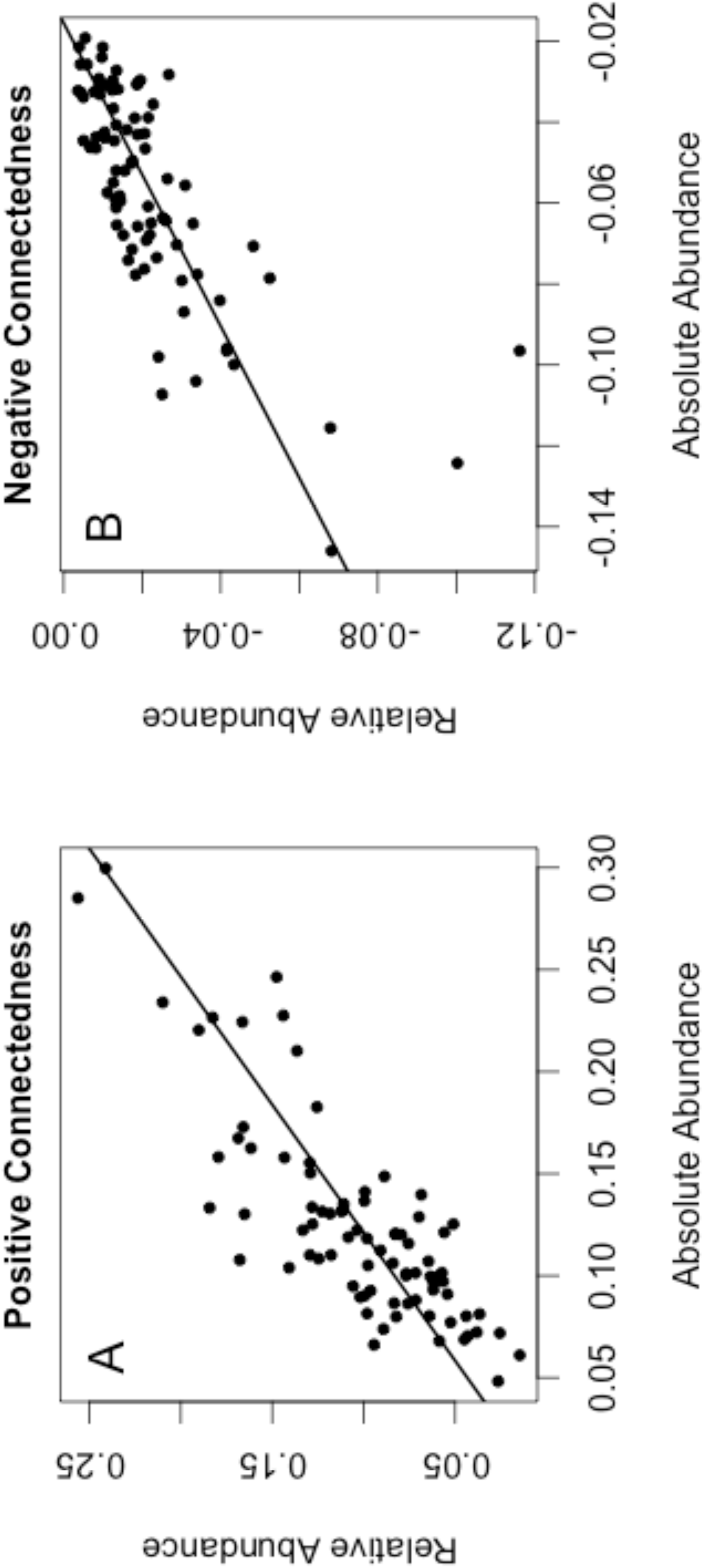
Comparing the metrics of connectedness obtained from the absolute abundance dataset (x-axes) and the relative abundance dataset (y-axes) shows agreement between the two methods of generating these metrics. Correlations between the metrics are 0.810 (panel A) and 0.741 (panel B). We used separate variables for positive and negative metrics because relativizing the dataset is expected to differentially affect positive and negative correlations. Solid lines show the fit of linear models.

#### 2. COHESION METRIC

##### Analysis

Many researchers seek to detect differences in community connectivity across time, space, or treatments. Thus, it would be useful to have a metric that quantifies, for each community, the degree to which its component taxa are connected. We used a simple algorithm to collapse the connectedness values of individual taxa into two parameters representing the connectivity of the entire sampled community, termed “cohesion.” Again, one metric of cohesion stems from positive correlations, and one metric stems from negative correlations. To calculate each cohesion metric, we multiplied the relative abundance of taxa in a sample by their associated connectedness values and summed these products. This cohesion index can be represented mathematically as the sum of the contribution of each of the *n* taxa in the community, after removing rare taxa (Eq. 1). Thus, communities with high abundances of strongly connected taxa would have a high score of community cohesion. We note that this index is bounded by −1 to 0 for negative cohesion or from 0 to 1 for positive cohesion.

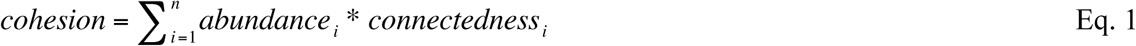

##### Validation

We had hypothesized that our cohesion metrics could be significant predictors of microbial community dynamics. Thus, a natural question to ask was whether our metrics of cohesion outperform environmental variables when analyzing the Lake Mendota data. Fortunately, the NTL LTER program has collected paired environmental data for the Lake Mendota phytoplankton samples. We obtained these environmental datasets to use as alternative predictors of community dynamics in Lake Mendota. The environmental datasets available (11 variables) were: water temperature, air temperature, dissolved oxygen concentration, dissolved oxygen saturation, Secchi depth, combined NO3 + NO2 concentrations, NH4 concentration, total nitrogen concentration, dissolved reactive phosphorus concentration, total phosphorus concentration, and dissolved silica concentrations. Protocols, data, and associated metadata can be found at https://lter.limnology.wisc.edu/. We use these environmental data to build an alternate model in our case study below.

#### 3. CASE STUDY OF UTILITY

##### Analysis

To demonstrate their utility, we applied our new metrics to the Lake Mendota phytoplankton dataset. We tested whether community cohesion could predict compositional turnover, a common response variable in microbial ecology. We used multiple regression to model compositional turnover (Bray-Curtis dissimilarity between time points) as a function of community cohesion at the initial time point. That is, Bray-Curis dissimilarity was the dependent variable, while positive and negative cohesion were the independent variables. Because time between samples influences Bray-Curtis dissimilarity (Nekola and White 1999, Shade et al. 2013), we only included pairs of samples taken within 36 to 48 days of each other. These criteria included 186 paired communities across the 19 years. Cohesion values (both positive and negative) were calculated at the first time point for each sample pair. We chose this timeframe because it was sufficiently long for multiple phytoplankton generations to have occurred, and because this timeframe was compatible with the sampling frequency.

Community cohesion was a strong predictor of compositional turnover (Fig. 4). The regression using our cohesion metrics explained 46.5% of variability (adjusted R^2^ = 0.465) in Bray-Curtis dissimilarity. Cohesion arising from negative correlations was a highly significant predictor, whereas cohesion arising from positive correlations was not significant (negative cohesion: F_1, 183_ = 6.81, p < 1* e ^-20^; positive cohesion: F_1, 183_ = 0.735, p = 0.405).

**Figure 4:**
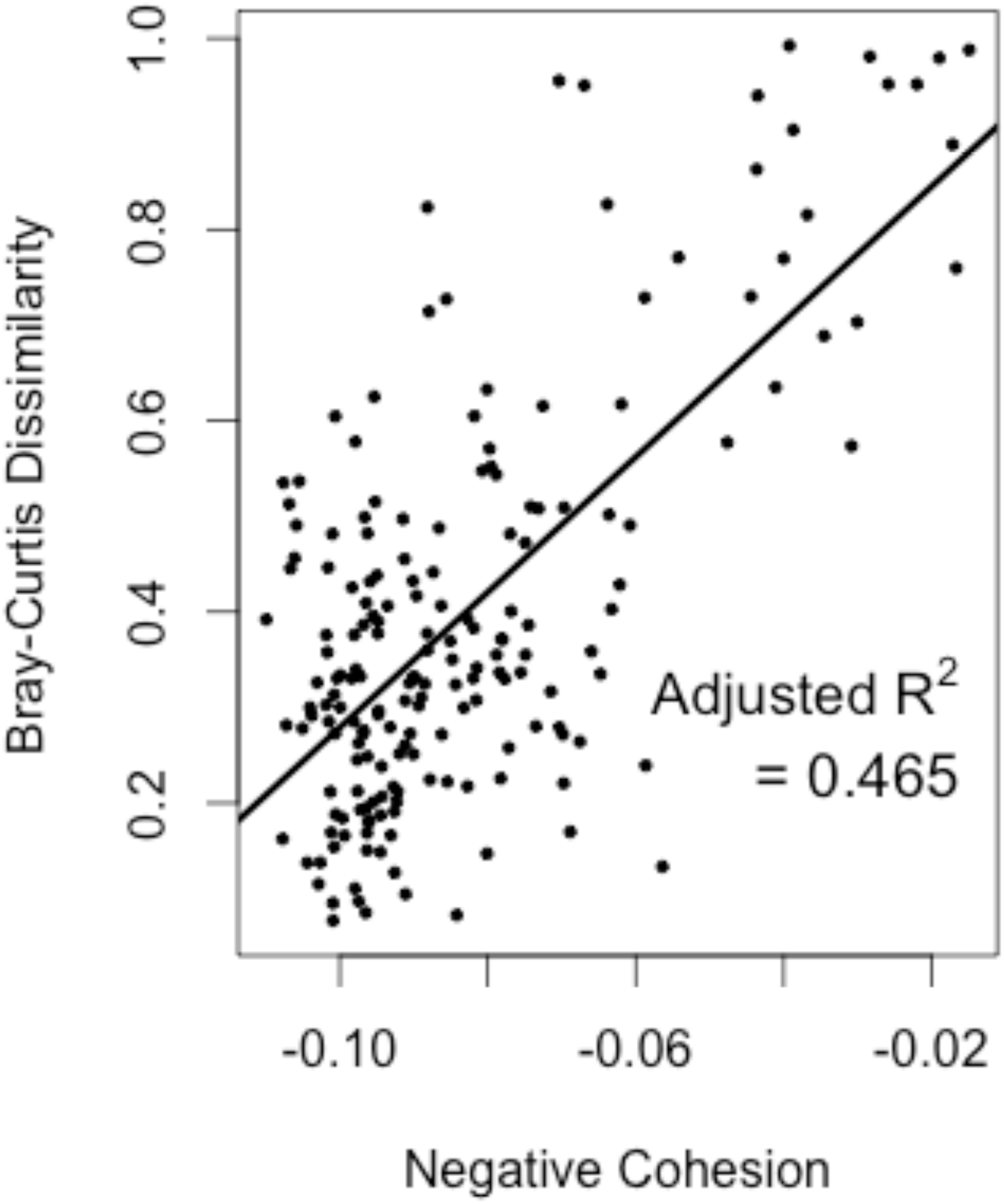
We used our metrics of community cohesion as predictors of the rate of compositional turnover (Bray-Curtis dissimilarity) in the Mendota phytoplankton communities. Negative cohesion was a significant predictor (p < 1*e^-20^) of Bray-Curtis dissimilarity, and the regression explained 46.5% of variation in compositional turnover.

For the purpose of model comparison, we used the associated environmental data to model Bray-Curtis dissimilarity as a function of environmental drivers. We included as predictors the 11 variables previously mentioned, as well as 11 additional predictors that measured the change in each of these variables between the two sample dates. Finally, because many chemical and biological processes are dependent on temperature (Brown et al. 2004), we included first order interactions between water temperature and the 21 other variables. We first included all 43 terms in the model, then used backward selection (which iteratively removes the least significant term in the model, beginning with interaction terms) until all remaining terms in the model were significant at p < 0.1, as to maximize the adjusted R^2^ value. Although this analysis does not represent an exhaustive list of possible environmental drivers, it includes all available paired environmental data from the long-term monitoring program. Twenty-nine values of Bray-Curtis dissimilarity were excluded from this analysis (leaving 157 of the 186 values), because they lacked one or more associated environmental variables. Additional details about this analysis can be found in the SOM.

In the final model after backward selection, 16 variables were retained as significant predictors (see SOM). The adjusted R^2^ of this model was 0.229. The *non-adjusted* R^2^ value of the full model (all 43 variables) was 0.393.

To address the generality of the relationship between cohesion and community turnover, we calculated cohesion metrics and Bray-Curtis dissimilarity for the four other phytoplankton datasets (Monona, Peter, Paul, and Tuesday lakes). Community cohesion was a significant predictor of Bray-Curtis dissimilarity in all datasets. In each instance, stronger cohesion resulting from negative correlations was related to lower compositional turnover. Table 1 presents the results of these analyses and associated workflow parameters. Additional information about the sensitivity of model performance to varying parameters can be found in the SOM.

**Table 1:**
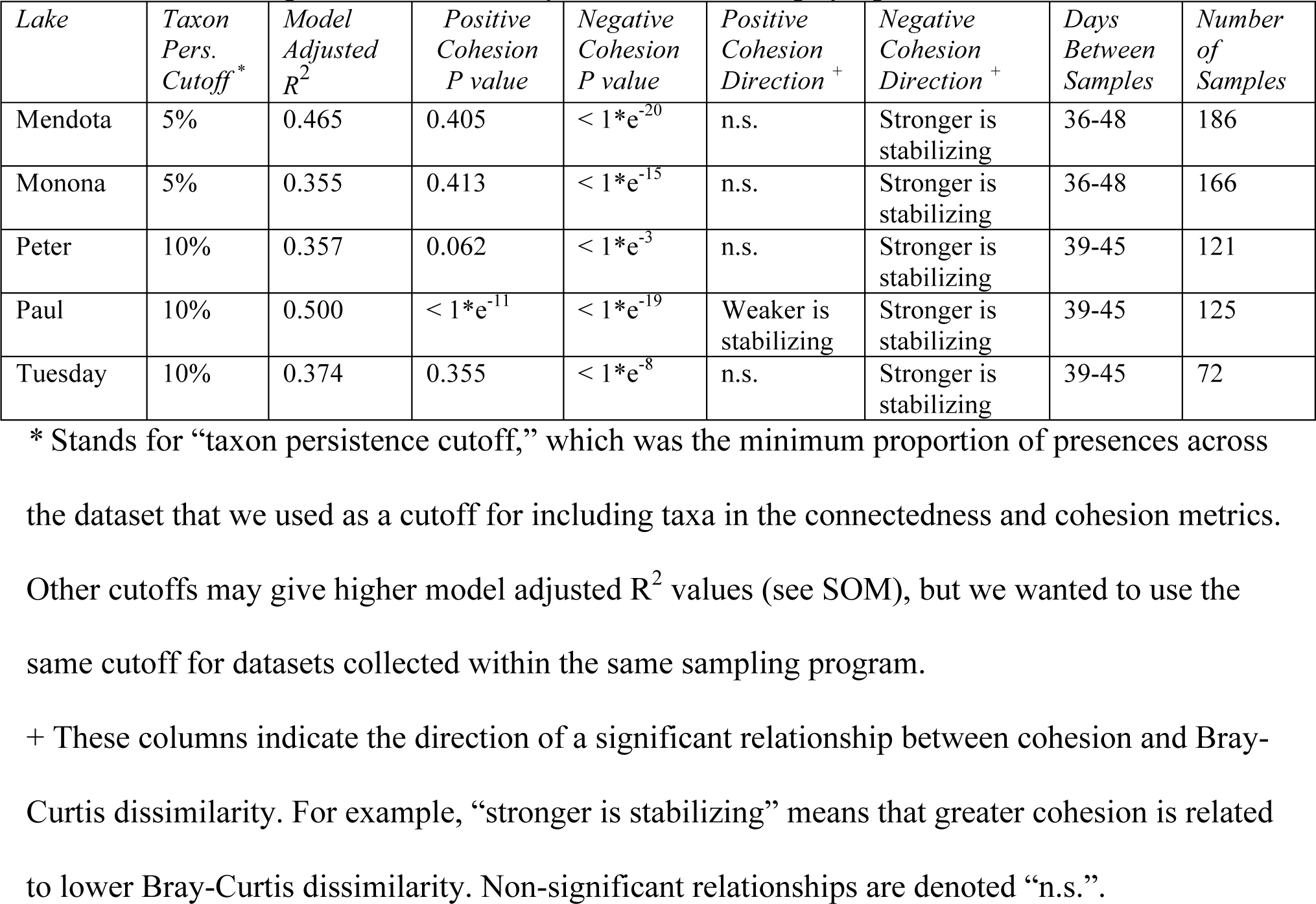
Cohesion predicts community turnover in five phytoplankton datasets

##### Validation

Strong correlations between predictor variables are known to influence the results of statistical analyses (Neter et al. 1996). Thus, we wondered whether strong correlations between taxa would necessarily generate the observed relationship where greater cohesion is related to lower compositional turnover. We conducted simulation studies to investigate whether our significant results might be simply an artifact of strong inter-taxon correlations. We generated datasets where taxa were highly correlated in abundance, as if they were synchronously responding to exogenous forces. We calculated cohesion metrics and Bray-Curtis dissimilarities for the simulated datasets to analyze whether strong taxon correlations was sufficient to produce results similar to those we observed in the real data.

Here, we briefly describe the process used to simulate datasets, while additional details can be found in SOM. First, we generated four autocorrelated vectors to represent exogenous forces, such as environmental drivers. Taxa were artificially correlated to these external vectors, thereby also producing strong correlations between taxa. We manipulated the taxon abundances to mimic other important features of the microbial datasets, including skewed taxon mean abundances and a large proportion of zeroes in the dataset. We calculated cohesion metrics and Bray-Curtis dissimilarities for the simulated datasets, and we used a multiple regression to model Bray-Curtis dissimilarity as a function of positive cohesion and negative cohesion. We recorded the R^2^ value and parameter estimates of this multiple regression. We repeated this simulation process 500 times to generate distributions of these results.

Our cohesion metrics had a very low ability to explain compositional turnover (Bray-Curtis dissimilarity) in the simulated datasets. The median model adjusted R^2^ value was 0.022, with 95% of adjusted R^2^ values below 0.088 (Fig. 5). Although the community cohesion metrics were highly significant predictors (p < 0.001) of community turnover more commonly than would be expected by chance (1.0% of simulations for positive cohesion and 8.6% for negative cohesion), the proportion of variance explained by these metrics was comparatively very low. For comparison, across the five phytoplankton datasets from Wisconsin lakes, model adjusted R^2^ values ranged from 0.36 to 0.50. Thus, there was comparatively little ability to explain compositional turnover in the simulated datasets using our cohesion metrics.

**Figure 5:**
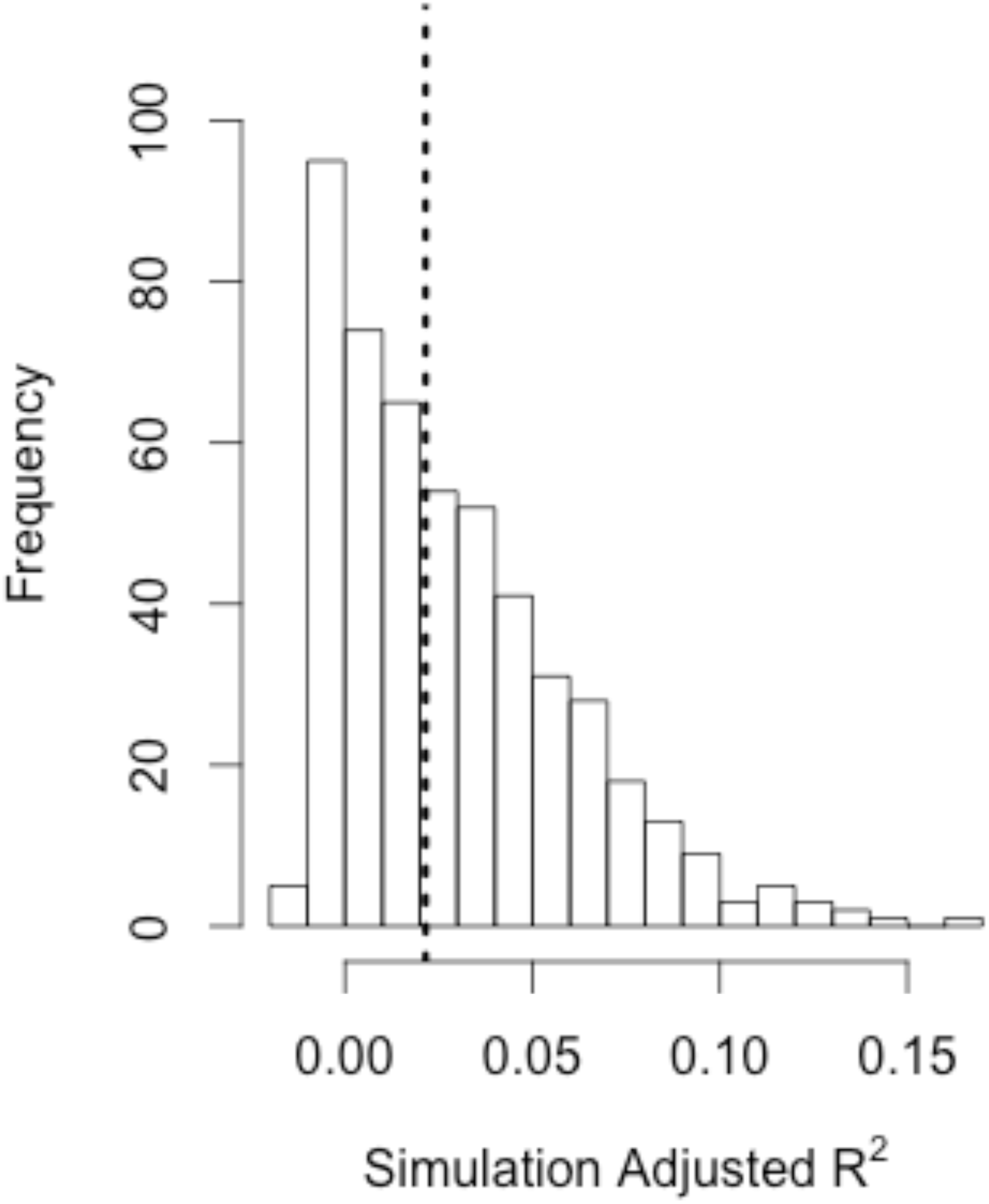
We simulated datasets where correlations between taxa were artificially produced by forced correlation to external factors. We calculated cohesion values for the simulated communities to test whether cohesion and Bray-Curtis dissimilarity were strongly related in simulated datasets. The histogram of model adjusted R^2^ values from our simulations shows that the median adjusted R^2^ was 0.022 (dashed line), with 95% of values falling below 0.088. For comparison, observed adjusted R^2^ values ranged from 0.36 to 0.50.

## Discussion

The ability to predict microbial community dynamics lags behind the amount of data collected in these systems (Blaser et al. 2016). Here, we present new metrics, called “cohesion,” which can be used as additional predictor variables in microbial models. The cohesion metrics contain information about the connectivity of microbial communities, which has been previously hypothesized to influence community dynamics (MacArthur 1955, May 1972, Nilsson and McCann 2016). Our cohesion metrics are easily calculated from a relative abundance table (R script provided online) and might be of interest to a variety of microbial ecologists and modelers.

In the Lake Mendota phytoplankton example, our two cohesion parameters outperformed the available environmental data at predicting phytoplankton community changes. The two cohesion parameters explained 46.5% of variability (adjusted R^2^ = 0.465) in community turnover over 19 years of phytoplankton sampling, in comparison to the final environmental model using 16 predictors, which explained 22.9% of community turnover (adjusted R^2^ = 0.229). Although there are almost certainly important predictors missing from the environmental model (e.g. photosynthetically active radiation, three-way interactions), the environmental model represents a commonly applied approach to explaining microbial compositional turnover (Tripathi et al. 2012, Chow et al. 2013) that uses all associated environmental data from a long-term sampling program. Although we still strongly advocate for the collection of environmental data, we note that cohesion was a much better predictor of compositional turnover than any available environmental variable.

Our workflow overcomes many challenges associated with using correlation-based techniques in microbial datasets. The validations we conducted indicated that our connectedness metrics are robust for relativized datasets, because connectedness metrics from relative and absolute datasets showed strong correspondence. Additionally, our cohesion metrics address a common problem of techniques describing community complexity (such as network analyses), which is that they do not quantify the connectivity of individual communities. For instance, the “hairball” generated from a network analysis has no parameters that are associated with each sample, and therefore the network cannot be used as a predictor variable. Thus, existing methods to quantify connectivity do not pair easily with other analyses. Our cohesion metrics quantify sample connectivity using only two parameters, which can be used as predictors in a variety of further analyses (linear regression, ordinations, time series, etc.). Finally, our simulations showed that strong inter-taxon correlations were not sufficient to reproduce the observed result that cohesion was a strong predictor of Bray-Curtis dissimilarity. In the simulations, cohesion had low explanatory power, even though taxa were highly correlated. From this result, we infer that correlations between taxa in real communities are an important aspect of complexity that is captured by our cohesion metrics.

Our cohesion metrics explain a significant amount of compositional change in all five phytoplankton datasets. Yet, it is not immediately clear what cohesion is measuring. There are two broad factors that could cause correlations between taxa: biotic interactions and environmental drivers. Thus, at least one of these two factors must underlie our connectedness and cohesion metrics. Here, we discuss the evidence supporting either of these interpretations:

### Cohesion as a Measure of Biotic Interactions

Even if shared responses to environmental drivers underlie most pairwise taxon correlations, cohesion could still indicate biotic interaction strength in a community. This would occur if taxa were influenced to the same degree by environmental drivers, but differentially influenced by species interactions. In this case, averaging over all correlations would give larger connectedness values for strong interactors and smaller connectedness values for weak interactors. Many studies have indicated that microbial taxa have differential interaction strengths. For example, some microbial communities contain keystone taxa, which have disproportionate effects on community dynamics through their strong taxon interactions (Trosvik and de Muinck 2015, Banerjee et al. 2016). Similarly, recent work suggests that many taxa within candidate phyla are obligate symbionts, meaning they must interact strongly with other taxa for their survival and reproduction (Kantor et al. 2013, Hug et al. 2016). Conversely, there are many taxa that can be modeled adequately as a function of environmental drivers; this is true for some bloom forming cyanobacteria, which are known to respond strongly to nutrient concentrations and temperature (McQueen and Lean 1987, Beaulieu et al. 2013). Taken together, these studies suggest that there is a wide spectrum of how strongly taxa interact with one another. These differences in interaction strength would be detected by our connectedness metric due to averaging over the large number of pairwise correlations. Thus, it is plausible that connectedness and cohesion are reflecting biotic interactions in communities.

We now examine our phytoplankton analysis results under the assumption that cohesion measures biotic interactions. The Bray-Curtis dissimilarity regression results would mean that communities with many strong interactors have lower rates of change, especially when the interactions create negative correlations between taxon abundances. This finding is in line with prior work showing that biotic interactions affect microbial community stability (Coyte et al. 2016). Thus, the interpretation that stronger biotic interactions lead to lower compositional turnover is a plausible explanation for our observed results. However, we specifically refrain from interpreting positive or negative connectedness values as indications of specific biotic interactions, such as predation, competition, or mutualism. For example, a positive correlation between two taxa could be the result of a mutualism between the taxa, or it could be the result of a shared predator declining in abundance. Further work, both empirical and theoretical, is necessary to identify what these positive and negative correlations signify in the context of the ecology of these organisms.

### Cohesion as a Measure of Environmental Synchrony

We now consider the possibility that connectedness and cohesion are simply detecting environmental synchrony. If a subset of taxa respond to a changing environmental driver, then these taxa will have strong pairwise correlations. For example, correlations between phytoplankton species of the same genus (and, therefore, with similar niches) can be upwards of 0.9, indicating strong similarity in abundance patterns. In this case, connectedness would measure the degree of environmentally-driven population synchrony that a taxon has with other taxa. A high cohesion value would indicate that a community has many taxa that respond simultaneously to external forces; then, cohesion would quantify overall community responsiveness to one or more environmental drivers. Under this assumption, cohesion should correlate with environmental drivers (e.g. cohesion is high because many taxa are positively correlated to warm temperatures, but cohesion drops when it gets colder and these taxa senesce). We tested this prediction with 22 variables from the environmental model (11 for the environmental variables and 11 for the changes in environmental variables) and found that negative cohesion in the Lake Mendota phytoplankton dataset generally had weak correlations with these predictors (absolute correlations < 0.25, SOM). We also looked for a seasonal trend in cohesion, but found no significant correlation between negative cohesion and Julian Day, or a quadratic term for Julian Day. Thus, we do not find any evidence that cohesion is simply reproducing the information contained in environmental data. Finally, our simulations show one example where taxon abundances could be driven exclusively by external factors (such as the environment), but this does not necessarily lead to strong predictive power of compositional turnover. However, our simulations omitted many features of real ecological communities, and so we cannot completely rule out the possibility that environmental drivers contributed to our cohesion metrics in the phytoplankton datasets.

Under the assumption that cohesion measures environmentally driven population synchrony, we examine our result that stronger cohesion was related to lower Bray-Curtis dissimilarity. In this scenario, communities that have strong cohesion contain high abundances of taxa that respond simultaneously to environmental forces. Then, communities with many synchronous taxa would turn over more slowly than communities with taxa whose abundances are independent of the environment. This conclusion is counterintuitive, but possible. This pattern could occur if taxa that are strongly influenced by the environment have lower variability than taxa that are weakly influenced by the environment; in that case, highly correlated taxa would have their abundances more tightly regulated than other taxa. Although plausible, this explanation disagrees with many studies that have found that environmental gradients regulate which taxa can persist in communities (Fierer and Jackson 2006, Walter and Ley 2011, Freedman and Zak 2015).

Comparing the two possible signals that cohesion might be detecting, we believe the evidence points to biotic interaction as the larger contributor. However, we expect that environmental synchrony is captured to some extent, with the relative importance of environmental factors depending on the particular communities and ecosystem. Regardless of the ecological force measured by cohesion, there is a clear result in the five datasets analyzed that stronger negative cohesion is related to lower compositional turnover. This result suggests that negative correlations between taxa are arranged non-randomly to counteract one another, thereby stabilizing community composition. In other words, relationships between taxa appear to form negative feedback loops that buffer changes in community composition. This result agrees with prior theoretical models that propose that feedback loops are integral to modulating food web stability (Neutel et al. 2007, Brose 2008). The finding that negative cohesion was stabilizing was not easily replicated in our simulations, where positive and negative correlations were interspersed with random magnitude throughout the dataset. Thus, the arrangement of correlations between taxa in the dataset appears to be an important feature of real communities that may contribute to their stability (Worm and Duffy 2003).

### Guidelines for Using Our Metrics

Although we used long-term time series datasets for the analyses presented here, our cohesion metrics can be used to predict community dynamics in a variety of datasets. For example, cohesion could be used with a spatially explicit dataset, where samples were collected from different locations across a landscape. In the context of phytoplankton samples, this could be a dataset consisting of samples from different locations in a lake or watershed. Then, the cohesion metrics could be used to predict community composition change at one location over time, or to predict differences in community composition between locations. It would also be interesting to investigate how cohesion is affected by experimental perturbations. Finally, cohesion could be used as a predictor in of many response variables. Additional applications of the cohesion metrics could include identifying communities susceptible to major compositional change (e.g. cyanobacterial blooms, infection in the human microbiome), relating community cohesion to spatial structure (e.g. how taxon connectedness relates to the dispersal abilities of different microbial taxa), and investigating how disturbance influences cohesion (e.g. how illness influences the cohesion of communities in a host-associated microbiome, how oil spills affect cohesion of marine microbial communities).

The critical step in the cohesion workflow is calculating reliable correlations between taxa. Thus, some datasets will be more suitable for our cohesion metric than others. For example, a dataset consisting of 20 samples from five lakes over multiple years might be a poor candidate for the cohesion metrics. In this case, correlations between taxa might be driven mainly by environmental differences or location, and the sample number would be too low to calculate robust correlations. Based on the phytoplankton datasets analyzed here, we suggest a lower limit of 40-50 samples when calculating cohesion metrics, with more samples necessary with more heterogeneous datasets. We also suggest including environmental variables as covariates when analyzing heterogeneous datasets. Finally, the persistence cutoff for including taxa should be adjusted based on the dataset being analyzed. For example, in datasets obtained by DNA sequencing, the sequencing depth affects taxon persistence (Smith and Peay 2014). Thus, it may be more effective to implement a cutoff by mean abundance.

## Conclusion

Our cohesion metrics provide a method to incorporate information about microbial community complexity into predictive models. These metrics are easy to calculate, needing only a relative abundance table. Furthermore, across all datasets analyzed in this study, cohesion was strongly related to compositional turnover. In systems where cohesion is a significant predictor of community properties (e.g. nutrient flux, rates of photosynthesis), this result could guide further investigation into the effects of microbial interactions in mediating community function. In this case, researchers might focus their efforts on understanding the role of highly connected taxa, which are identified in our workflow. We aim to eventually determine the features that distinguish systems in which cohesion is important versus systems in which cohesion does not predict community properties.

## Acknowledgements

We thank the North Temperate Lakes LTER program for the use of their publicly available data on Lake Mendota and Lake Monona. We also thank the Cascade research group for the use of their data from Peter, Paul, and Tuesday lakes, which is hosted on the LTER website. This manuscript has been much improved as a result of comments from the McMahon lab. This work was funded by a United States National Science Foundation (NSF) GRFP award to CMH (DGE-1256259). KDM acknowledges funding from the NSF Long Term Ecological Research program (NTL-LTER DEB-1440297) and an INSPIRE award (DEB-1344254).

Supplementary material is available at ISME Journal’s website.

